# Mutant *KRAS* dosage contributes to heterogeneity in lung cancer therapeutic response

**DOI:** 10.64898/2026.05.06.723208

**Authors:** Alice T. Browne, Chris McCann, William J. McDaid, Nikita Lewis, Sandhya Sridhar, Gary Doherty, Deborah Y. Moss, Matilda Downs, Stephen Marry, Aaron Phillips, Connor N. Brown, Alyson Speed, Gemma Logan, Gera Jellema, James Bradford, Catherine Davidson, Vicky Coyle, Donna Small, Nick Orr, Richard Kennedy, Sarah Maguire, Carla P. Martins, Emma M. Kerr

## Abstract

Oncogenic *KRAS* mutations promote tumorigenesis by constitutive activation of multiple, well-characterised signalling pathways. However, there is significant heterogeneity across mutant *KRAS* tumours in terms of mutation present, mutant allele abundance and downstream signalling strength. It is unclear whether these variations can impact responses to specific therapies. Here, we demonstrate that ∼20% of lung adenocarcinomas (LUAD) show an increase in mutant KRAS dosage (KRAS^mutant^ allele fraction > KRAS^wild-type^). Furthermore, we show that *KRAS* mutant dosage can directly influence clinical outcome and therapeutic susceptibilities in lung cancer. Our findings show that mutant *KRAS* copy gains specifically affect platinum lung cancer response, promoting resistance to this standard-of-care therapy. Importantly, increases in *KRAS* mutant dosage are also associated with an increased vulnerability to pS6K inhibition, due to the unique metabolic rewiring of these cells. Together, we show that mutant *KRAS* dosage contributes to the phenotypic heterogeneity of mutant KRAS NSCLC and that assessment of mutant KRAS content or signalling strength can help optimise treatments strategies for these patients.

## Introduction

Lung cancer remains the most lethal cancer worldwide, with high mortality rates due to frequent late diagnosis, a paucity of treatment options, and heterogeneous treatment responses. Oncogenic *KRAS* mutations are found in ∼30% of lung adenocarcinomas^1^ and can have a myriad of effects on tumour cells and the tumour microenvironment (TME), influencing proliferation, invasion, metabolism and inflammation^2,3^. KRAS mutations have historically limited treatment options for patients given the paucity of direct inhibitors^4^ and lack of response to other targeted therapies. Therefore, until recently, chemotherapy and radiotherapy combinations were the most common treatment options for *KRAS* mutant lung cancer patients, but responses remain variable and hard to predict. More recently, immunotherapy and direct KRAS^G12C^-inhibitors have been added to the lung cancer therapy arsenal, opening a promising new chapter in the treatment of these cancers. However, even in the case of KRAS^G12C^-inhibitors, multiple mechanisms of resistance were already reported^5^, highlighting the complexity in effectively treating this group of cancers.

Part of the phenotypic heterogeneity of mutant KRAS NSCLC may be contributed by KRAS mutant status itself, which varies significantly between tumours. Accordingly, the type of mutation (e.g. KRAS^G12D^ vs KRAS^G12V^) and mutant allele/protein abundance relative to KRAS wild-type (allelic balance) can have a significant impact on tumour signalling, fitness and potentially, clinical outcome. In agreement, we previously reported that heterogeneity in mutant *Kras* gene copy number is common within genetically engineered mouse models of lung adenocarcinoma (LUAD), with mutant-specific copy gains (i.e. amplifications +/- WT loss) being selected for during progression to advanced disease^6^. These *Kras* mutant-specific copy gains, which are enriched in advanced lung tumours, drive the rewiring of glucose utilisation towards glycolysis/TCA cycle metabolism and redox buffering through glutathione production and ultimately result in a therapeutically-exploitable dependency on glucose/glutathione metabolism^7^. Mutant *KRAS* copy gains (*KRAS MutCG*) are also prevalent in human LUAD^7–9^, but the clinical implication of these alterations is unknown. Interestingly, *KRAS* mutant allele specific amplifications have recently been associated with distinct therapeutic susceptibilities such as increased MEK inhibitor sensitivity^9,10^ and acquired resistance to KRAS^G12C^-inhibitors^11,12^, indicating that mutant KRAS copy content can influence therapeutic responses. A broader, in-depth understanding of the impact of mutant KRAS copy content on LUAD-treatment response could help improve clinical outcomes to current and future therapies.

Using combined *in silico*, *in vitro* and *in vivo* approaches, we now show that *KRAS MutCG* is associated with worse clinical outcome in NSCLC, relative to copy-neutral KRAS mutant lung cancers (KRAS mutant cancers with no evidence of mutant specific copy gain) and promote intrinsic resistance to distinct DNA-damaging chemotherapies. Furthermore, we demonstrated that this decreased sensitivity to specific chemotherapies can be predicted based on allelic profiling (*MutCG* vs copy-neutral) or *MutCG*-driven signalling. Importantly, we also show that the unique molecular signatures of this aggressive subset of *KRAS* mutant lung cancers can also be exploited therapeutically. In particular, our analyses uncovered a novel dependency of *KRAS MutCG* tumour cells on p70 S6 Kinase (p70S6K)-S6 signalling, which is directly linked to MutCG-driven reprogramming of glucose metabolism.

## Results

### KRAS Mut Copy Gain drive resistance to platinum-based therapy

Platinum doublets have been the backbone of chemotherapy treatment for lung cancer patients with advanced disease^13^, including mutant KRAS cancers. Platinum drugs, and cisplatin in particular, are well known to induce metabolic stress via induction of reactive oxygen species (ROS)^14^, a stress that KRAS MutCG cell models were shown to more readily detoxify^7^. Given this, we started by asking whether *KRAS MutCG* could impact the sensitivity to platinum chemotherapy treatment in LUAD patients. We first examined the prevalence of *KRAS^mutant^*allele specific copy gains in human LUAD samples in cBioportal (LUAD PanCancer ^15–17^) by analysing *KRAS^mutant^* DNA variant allele frequency (VAF), defined per sample as variant *KRAS^mutant^* reads/ total *KRAS* reads. Substantial heterogeneity is observed in the *KRAS^mutant^* allele fraction, ranging from 0.08 to 0.95, with a mean of 0.37 ± 0.18 s.d. (Figure 1a). Importantly, of the 168 samples assessed, one fifth (20.24%) display *KRAS^mutant^* VAF of >0.5, which we hereafter define as “Mutant Copy Gain” (MutCG, Figure 1b), with the *KRAS^mutant^* VAF 0-0.49 group termed “Mutant Copy Neutral” (MutCN, Figure 1b). This PanCancer cohort contains detailed treatment information and includes forty-five patients with *KRAS^mutant^* LUAD that received cisplatin or carboplatin treatment. Despite the small numbers available for survival analysis, we found that in this study patients with MutCG (n=12) had nearly three times shorter median overall survival relative to patients in the Mut group (Figure 1c, MutCG 27.16 months, MutCN 78.67 months, p=0.0641, hazard ratio MutCG/MutCN: 2.493). Importantly, this cohort also showed a direct correlation between increased mutant allele content and enhanced KRAS expression at mRNA level (Extended data Figure 1a); as well as with increased glycolytic gene expression, assessed using two independent Glycolysis scores: MSigDB Hallmark Glycolysis^18^ (Figure 1d, Ext. data Figure 1b) and the Glycolysis Related score^19^ (Figure 1e, Ext. data Figure 1c).

**Figure 1:**
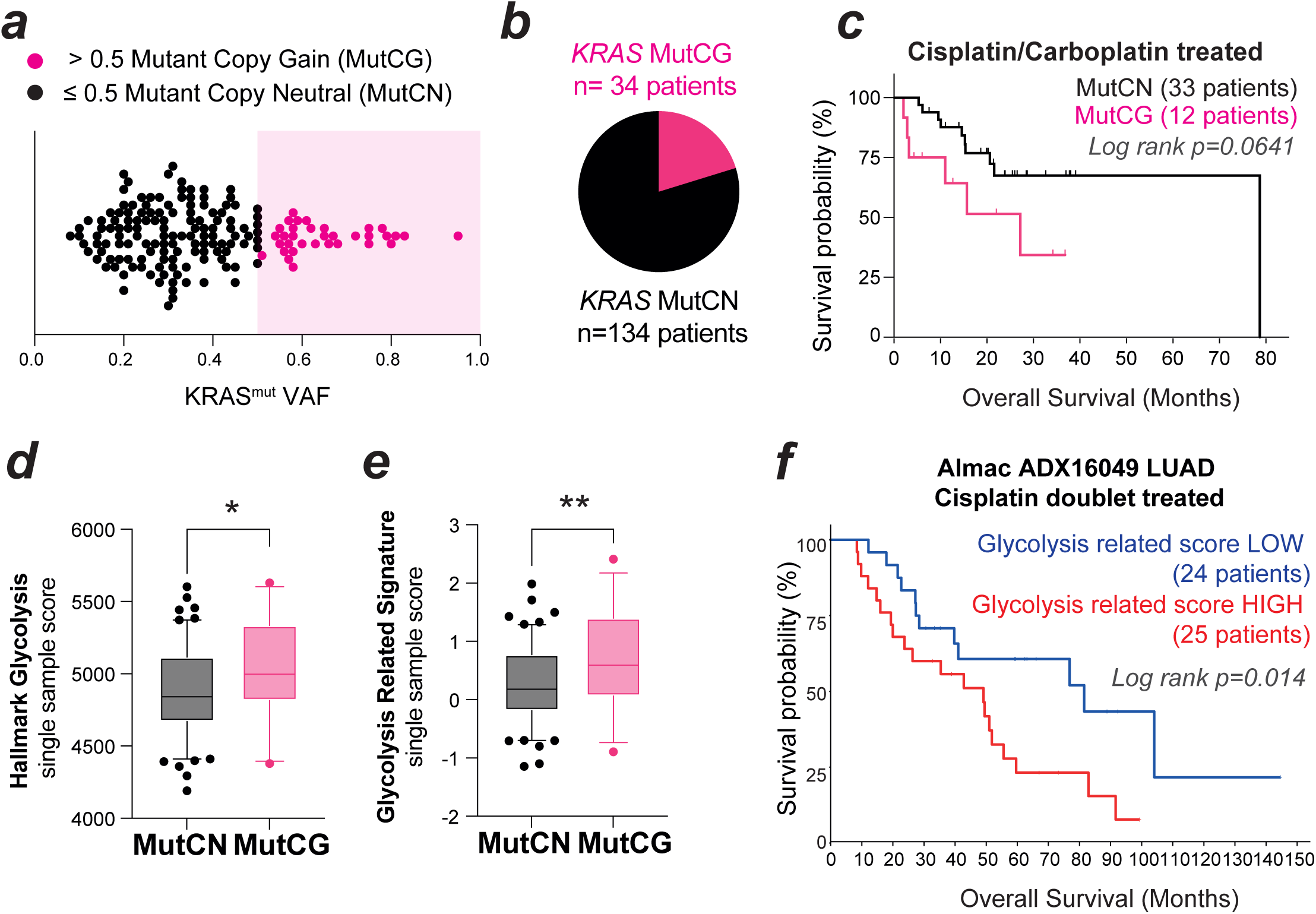
*KRAS^mutant^* copy gain driven metabolic reprogramming predicts chemoresistance in Lung Adenocarcinoma. Reference and Variant read counts across the *KRAS* locus for *KRAS^mutant^* TCGA PanCancer lung adenocarcinoma samples are used to determine *KRAS* mutant Variant Allele Frequency (VAF) per sample (a), and identify patients with Mutant Copy Gain (MutCG, b) defined as VAF>0.5 or Mutant Copy Neutral (MutCN, b) defined as VAF<0.5; (c) Overall survival analysis of Cisplatin/Carboplatin treated patients classified as MutCN (black, n=33) or MutCG (pink, n=12), with log-rank test p=0.0641; single sample scores for MutCN and MutCG groups for MSigDB Hallmark Glycolysis (d) or Glycolysis Related Signature (e, Liu et al, 2019), box plots show median with 25th to 75th percentiles, whiskers represent 5th and 95th percentiles. Pairwise t-test *p<0.05, **p<0.01; Overall survival analysis of Cisplatin/Vinorelbine treated ADX16049 Lung Adenocarcinoma patients classified as Glycolysis Score Low (blue, n=24) or Glycolysis Score high (red, n=25)

To further validate these findings, we accessed additional locally available cohorts. As VAF and treatment information were not readily available within the same datasets/patient cohorts, we used transcriptional changes in glycolytic gene expression as a surrogate for MutCG, since we previously showed that these changes correlate directly with MutCG^7^. Analyses of a second cohort of LUAD patients treated with cisplatin/vinorelbine doublet chemotherapy (ADX16049, n=49 patients) showed that tumours with High Glycolysis Related scores also display platinum-resistance, as indicated by their shorter overall survival (Figure 1f, p=0.014). Together, these analyses show that KRAS MutCG in human LUAD is associated with reprogramming of glucose metabolism^7^, and correlates with intrinsic resistance to platinum-based chemotherapy.

To characterise the mechanistic basis for increased resistance to platinum in MutCG tumours we tested platinum-responses in a panel of eight *KRAS^mutant^*non-small cell lung cancer (NSCLC) cell lines *in vitro*. These cell lines carried different *KRAS* hotspot mutations, but the majority (5/8) had G12C alterations (Figure 2a). KRAS zygosity status was attributed according to COSMIC annotation (presence/absence of wild-type; WT) and was further validated by Sanger sequencing. Accordingly, 4 cell lines were classified as mutant heterozygous (HET) and 4 as mutant homozygous (HOM). The relative mutant content of each cell line (variant allele frequency; VAF) was also determined by pyrosequencing and ranged from 0.4-1.0, (HET mean= 0.56; HOM mean= 0.98, Ext. data Figure 2a), in line with previously reported VAF for these models^20^. We confirmed that the reported *MutCG*-associated phenotypes were recapitulated in this new cell panel. Namely, as previously reported for other cellular models^7^, HOM cells exhibited significantly enhanced RAS activation, glycolysis (indicated by ECAR) and sensitivity to 2DG, relative to HET cells (Ext. data Figure 2). HOM cells also displayed increased colony forming potential, indicative of increased aggressiveness, despite showing no significant difference in growth rate. Reassuringly, in the current panel, increased RAS activity, glycolytic flux and 2DG sensitivity directly correlated with increased VAF (Figure 2b-d). Noteworthy, a range of other mutations reported in *KRAS^mutant^*lung cancer were also represented in the cell panel, such as alterations in *TP53*, *STK11* and *CDKN2A* (Figure 2a), but none of the phenotypes tested correlated with the presence/absence of *TP53* or *STK11* status (Ext. data Figure 2g-h).

**Figure 2:**
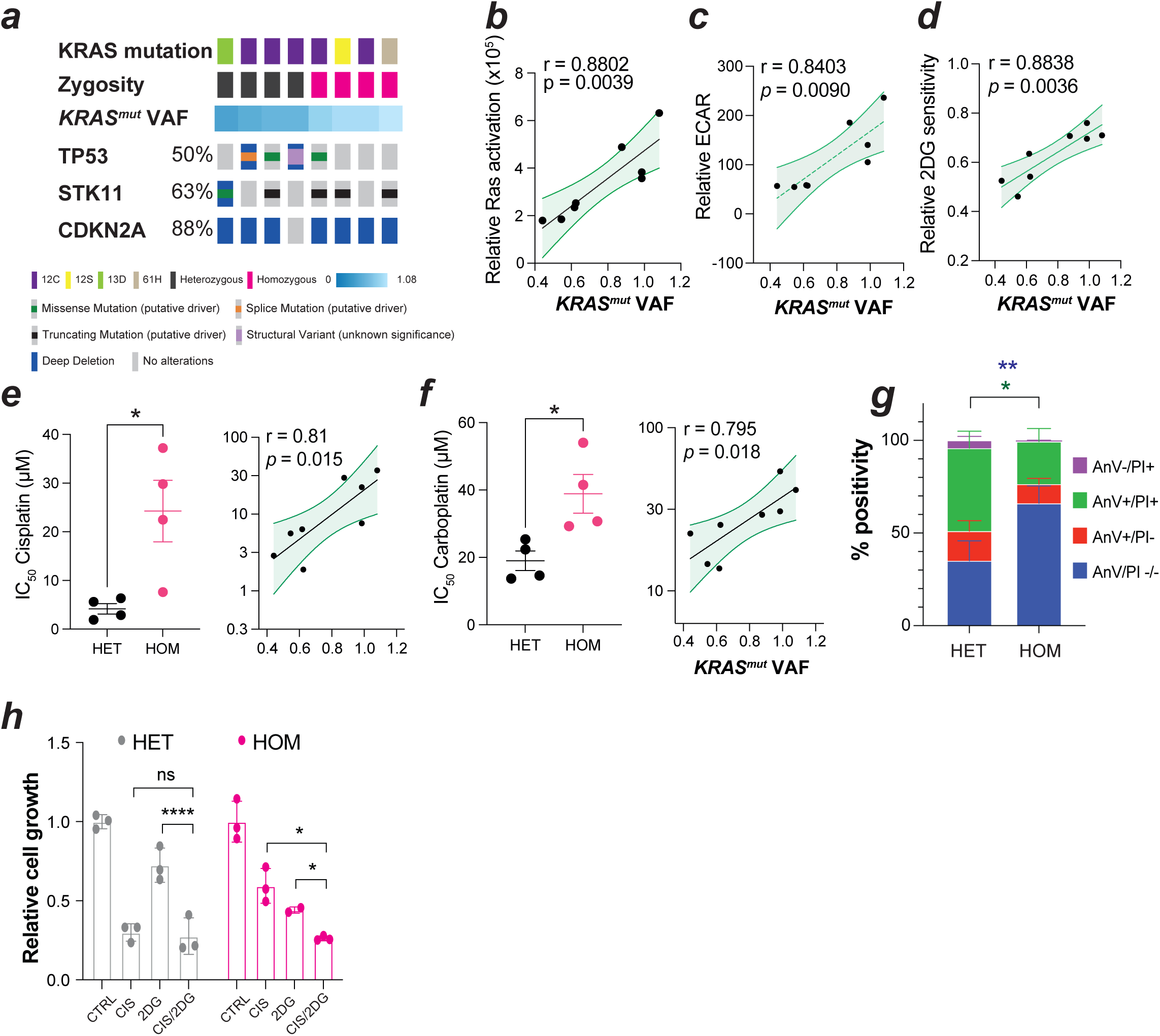
*KRAS^mutant^* copy gains promote resistance to platinum chemotherapy. (a) Summary illustration of KRAS mutation, zygosity, VAF status as well as alterations in TP53, STK11 and CDKN2A in 8 NSCLC cell lines; Correlation of relative Ras activity (b), Extracellular Acidification Rate (ECAR, c) and 2DG sensitivity (d) to KRAS mutant VAF; IC_50_ values for Cisplatin (e) and Carboplatin (f) in cell line panel, presented by zygosity status (HETerozygous/HOMozygous) or correlated with KRAS mutant VAF; (g) Annexin V/PI positivity in HET and HOM cell lines treated with cisplatin for 72hrs; (h) relative cell growth in response to cisplatin, 2DG or combination treatment for 72hrs determined by CV staining. Datapoints represent individual cell line mean of technical replicates (b-h) and line shows genotype mean ±sem (e-g) or linear regression analysis ±95% confidence interval (b-f) with p and r values determined by t-test (e,f) two-way ANOVA with mutiple testing (g, h) or Pearson’s correlation analysis (b-f) as appropriate; Significance indicated, *p<0.05, **p<0.01, ****p<0.0001, ns = not significant.

To directly assess the potential impact of MutCG on the response to platinum, HOM and HET cells were treated for 72 hrs with cisplatin and carboplatin and the concentration that caused 50% reduction in viability (IC_50_) determined by cell titre glo. Consistent with our observations in the patient cohorts,-HOM NSCLC cells were significantly more resistant to cisplatin and, albeit to a lesser extent, carboplatin, than HET cells. Accordingly, HOM cells showed a 5.8-fold (cisplatin) and 2-fold (carboplatin) increase in mean GIC_50_ relative to HET cells (Figure 2e-f), with sensitivity to platinum agents directly correlated to *KRAS^mutant^* dose (Figure 2e-f, *KRAS^mut^* VAF). To independently confirm this differential sensitivity we assessed the impact of cisplatin and carboplatin by ortholog assays, namely short-term (cell number) and long-term (colony formation) growth rates analysis (Ext. data Figure 3a-c) and induction of cell death (Figure 2g) in the 8 cell line panel and also, in a previously characterised murine matched HET/HOM Kras^G12D^;p53^null^ cell line pair^7^ (mHET and mHOM; Ext. data Figure 3d). These analyses confirmed that HET cells were significantly more sensitive to cisplatin treatment, displaying decreased growth and higher levels of cell death upon platinum treatment, relative to HOM cells.

We next asked if the decreased sensitivity of *KRAS MutCG* cells to platinum-based chemotherapy was reflective of a multi-drug resistance phenotype. To address this, we compared the sensitivity of the panel to a broad range of standard-of-care chemo- and targeted-therapies used in NSCLC, as well as agents targeting RAS-dependent TKI signalling pathways. The compounds examined included multiple DNA-damaging chemotherapies such as taxanes (docetaxel, paclitaxel), vinca alkaloids (vinorelbine), antifolates (pemetrexed) and anthracyclines (doxorubicin), as well as targeted TKI agents (erlotinib, selumetinib, neratinib, afatinib). Interestingly, no significant difference in sensitivity was observed between HET/HOM cells against these agents (Ext. data figure 3e). In agreement, no significant correlation was observed between IC_50_ values and *KRAS^mutant^* dose (Ext. data figure 3f) for any of these compounds. Taken together, these data demonstrate that *KRAS^mutant^* content has a specific impact on platinum-sensitivity in lung cancer.

Lastly, to determine whether the enhanced glucose metabolism of HOM cells contributed to their platinum resistance we co-treated HET and HOM cells with cisplatin and the glycolysis inhibitor 2DG for 72hrs and determined its impact on growth. Interestingly, while 2DG alone reduced the growth of both HOM and HET cells, 2DG-mediated glycolysis inhibition did not impact the response of HET cells to cisplatin. In contrast 2DG treatment significantly impacted cisplatin response in HOM cells, which in the presence of 2DG became as responsive to cisplatin as HET cells (Figure 2h). The (re)sensitisation of HOM cells to cisplatin through glycolysis inhibition indicates that *KRAS MutCG*-driven metabolic rewiring contributes directly to their resistance to platinum.

The main cytotoxic and cytostatic effects of cisplatin are thought to derive from its binding to DNA, with the resulting cisplatin/DNA adducts acting as potent activators of the DNA damage response^21,22^. Cisplatin exposure can also increase reactive oxygen species (ROS), which subsequently can induce DNA damage in cells^14,23,24^. A well described mechanism of cisplatin detoxification is through the action of glutathione (GSH)^25,26^. The formation of GSH/cisplatin adducts depletes the free pool-of cisplatin in cells, and can therefore contribute to resistance to treatment. We previously showed that GSH levels were upregulated in HOM cells and that this increase was directly linked to glucose metabolism rewiring driven by enhanced KRAS^mutant^ activity^7^. The increased resistance of HOM cells to cisplatin suggests that *KRAS MutCG* can drive resistance to other ROS-inducing DNA damaging agents, such as ionising radiation (IR), another common lung cancer therapy ^27^. In agreement, HOM cells were significantly more resistant to IR treatment than HET cells (Ext. data figure 3g-h). These data indicate that *KRAS* MutCG-driven GSH-upregulation may contribute to resistance in the clinic to ROS-inducing agents.

Collectively, these data comprehensively show that *KRAS^mutant^* dosage can impact clinical responses to platinum-based chemotherapy and radiotherapy and indicates that *KRAS* VAF and metabolic signatures could be useful clinical biomarkers in this setting.

### Mutant KRAS dosage influences sensitivity to targeted therapies

Given that the unique molecular signatures associated with MutCG can drive resistance to some therapeutic agents, we next hypothesised that this aggressive group of lung cancers may conversely display enhanced sensitivity to other treatments, namely targeted therapies. Using publicly available drug IC_50_ screening data from the Genomics of Drug Sensitivity in Cancer (GDSC) project^28^ we examined sensitivity to 99 compounds across 29 *KRAS^mutant^* NSCLC cell lines. KRAS^mutant^ content was stratified based on zygosity (HET v HOM) or *KRAS^mutant^* VAF. As shown in Figure 3a, this analysis identified a subset of compounds that displayed increased potency in HOM cells over HET, and this potency correlated strongly with MutCG (lower left quadrant).

**Figure 3:**
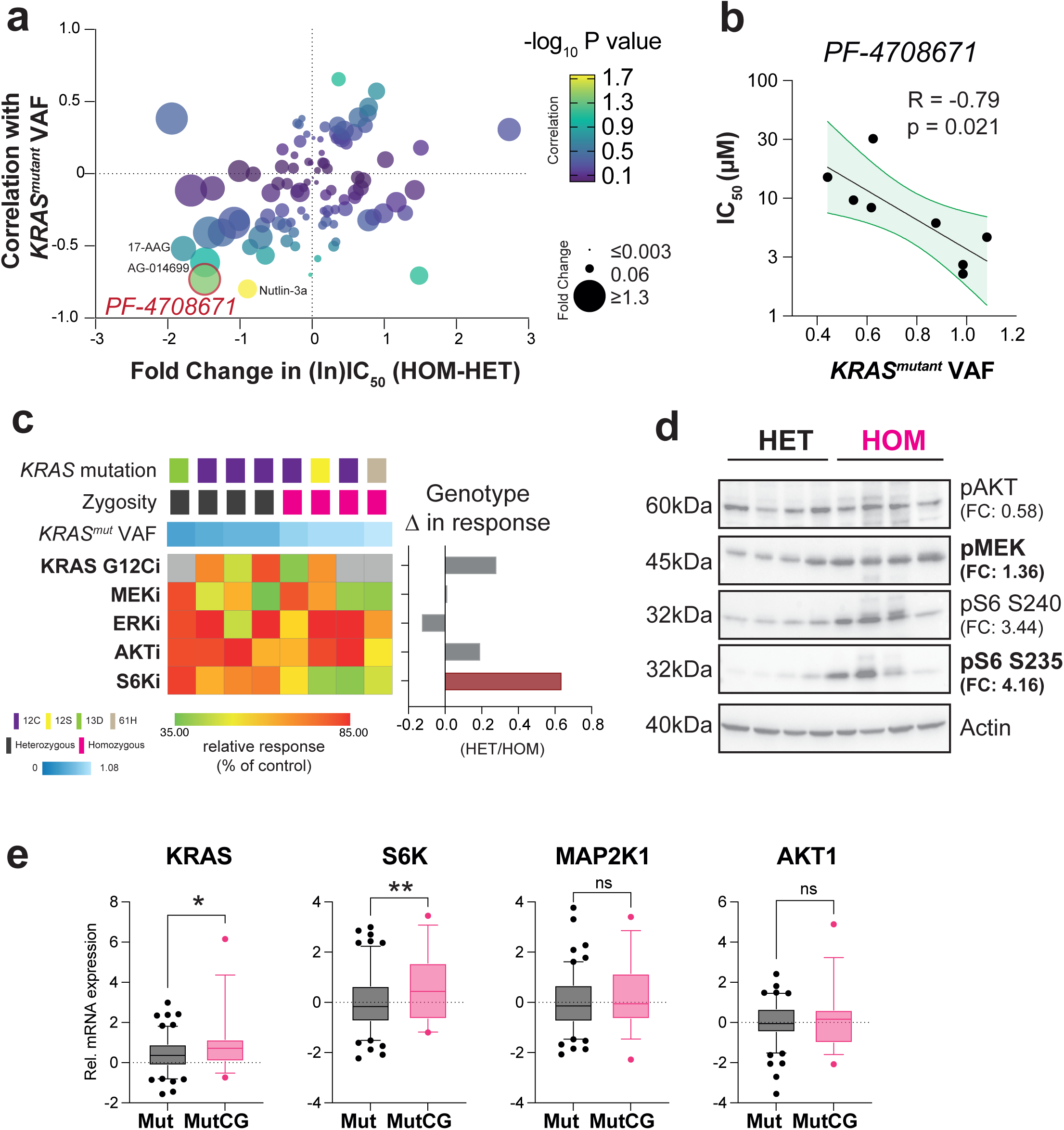
Analysis of drug senstivities highlights differential signalling axis with *KRAS^mutant^* copy gain. (a) Genomics of Drug Sensitivity IC_50_ data for *KRAS* mutant NSCLC cell lines was analysed to determine impact of MutCG, comparing genotype effect by t-test (fold change on x-axis) and mutant doseage effect by Pearson’s correlation (R value shown on y-axis). P values are indicated by spot size (t-test) and colour (correlation). Location of PF-4708671 is highlighted. (b) Validation of direct correlation between senstivity to PF-4708671 and MutCG in panel of 8 NSCLC cell lines, with Pearson’s R and P value indicated. Dots represent IC_50_ of individual cell lines, with linear regression line ± 95% confidence interval shown; (c) Sensitivity to KRAS pathway inhibitors, presented as % of vehicle control per cell line, in 8 cell line panel, with KRAS mutation, zygosity and VAF indicated. Change in response relative to vehicle control per cell line, and genotype difference calculated as mean HET/mean HOM values, pairwise comparison between genotypes analysed by t-test, significant difference indicated by red colour bar, adjP<0.05; (d) Relative protein expression in 8 NSCLC lines, analysed by t-test following densitometry normalised to Actin control, with genotype difference HOM/HET fold change shown and p value<0.05 indicated by bold text; (e) relative expression represented by mRNA z-scores per patient from TCGA Firehose cohort. Box plots show median with 25th to 75th percentiles, whiskers represent 5th and 95th percentiles, analysed by t-test, *p<0.05, **p<0.01.

PF-4708671, an inhibitor of p70 S6 Kinase (p70S6K) ^29^ showed the strongest MutCG-selectivity: 2.8 fold difference in mean IC_50_ HOM v HET, Pearson’s correlation with *KRAS^mutant^*VAF R = -0.7319, p = 0.039 (Figure 3a). We independently verified this MutCG-selectivity in our 8 cell line panel. As shown, IC_50_ of PF-4708671 directly correlated with *KRAS^mutant^*VAF (Figure 3b, Pearson’s R = -0.79, p = 0.021). We also verified other hits for MutCG-selectivity in our 8 cell line panel, namely 17-AAG and AG-014699 (Ext. data figure 4a-b). These showed the same direct correlation with *KRAS^mutant^* VAF.

p70S6K is a serine/threonine kinase directly responsible for phosphorylating S6, a well-established target of mTOR signalling^30,31^ and co-ordinator of KRAS-driven metabolic reprogramming ^32,33^. Therefore, we next asked if a similar MutGC selectivity could be observed with KRAS inhibition directly, or with other KRAS-pathway targeting agents, by testing the impact of KRAS loss or inhibition by G12C inhibitors (G12Ci-12, MRTX859), as well as inhibitors of MEK (PD-0325901), ERK (SCH772984), and AKT (MK-2206) inhibitors in our cell panel, relative to PF-4708671.

KRAS dependency in KRAS mutant NSCLC cell models was first explored using DepMap data^34^ (https://depmap.org/portal). CRISPR gene effect scores were comparable between HET and HOM lines (Ext. data figure 5a) and showed similar sensitivity to the KRAS G12C inhibitor (G12Ci) (Ext. data figure 5b), suggesting that all mutant cell lines require continued KRAS signalling. We confirmed these findings by testing MRTX859 sensitivity in the subset of 5 G12C cell lines in our panel (Ext. data figure 5c). Pairwise comparison of HET/HOM response demonstrated that there was no difference in sensitivity to direct KRAS G12C inhibition under the conditions tested (Figure 3c).

Intriguingly, of all the RAS-pathway targeting compounds tested, only PF-4708671 was significantly differentially effective between the HOM and HET genotypes (red, Figure 3c). These data suggests that MutCG may promote discrete differences in signalling within the RAS pathway, rather than a generalised increase in MAPK activity. To test that, we analysed RAS pathway activation by Western blot, to determine if phosphorylation levels of MEK, AKT and S6 (Figure 3d) were impacted by MutCG. We observed no difference in pAKT, a modest increase in pMEK (FC:1.36) and a striking increase in pS6 (FC:4.16 S235) in HOM cells relative to HET cells, supporting this distinct sensitivity to PF-4708671. Lastly, we asked if similar differences could be observed in patient samples. Given the scarcity of protein data, we analysed the differential expression of RAS pathway transcripts in TCGA cohorts of *KRAS* Mut (i.e copy number neutral) and MutCG samples (defined in Figure 1b). Significant increases in KRAS and S6K mRNA were observed in MutCG samples, but expression of MAP2K1 and AKT1 remained comparable between the two groups (Figure 3e). Taken together, these data suggest that *KRAS MutCG* promotes a significant and specific increase in p70S6K/S6 signalling, which results in a potentially targetable dependency.

### KRAS^mutant^ copy gains drive an increased dependency on S6K signalling in lung cancer

We next asked if the sensitivity of HOM cells to S6K-inhibition could be observed in vivo. We started by testing the impact of PF-4708671 on the colony-formation potential of HET (H23) and HOM (H2030) cells. As previously shown, HOMs cells form colonies more rapidly than HET cells (Ext. data figure 2d), highlighting their more aggressive phenotype. Interestingly, PF-4708671 significantly inhibited colony growth of HOM cells, but had a milder effect in HET cells (Figure 4a), indicating that S6K activity plays an important role on the invasive phenotype of HOM cells. We then tested this phenotype in vivo using subcutaneous xenograft transplantation of the same HET and HOM cells. Given that HOM cells grow more rapidly than HET cells in vivo^7^, in both cases daily treatment with PF-4708671 (50mg/kg via i.p.) or vehicle was initiated when tumours volume reached 100mm^3^. HOM tumours showed significant growth inhibition after 3 and 6 days of PF-4708671 treatment, while no significant difference in tumour volume was observed between vehicle and PF-4708671 -treated HET tumours (Figure 4b).

**Figure 4:**
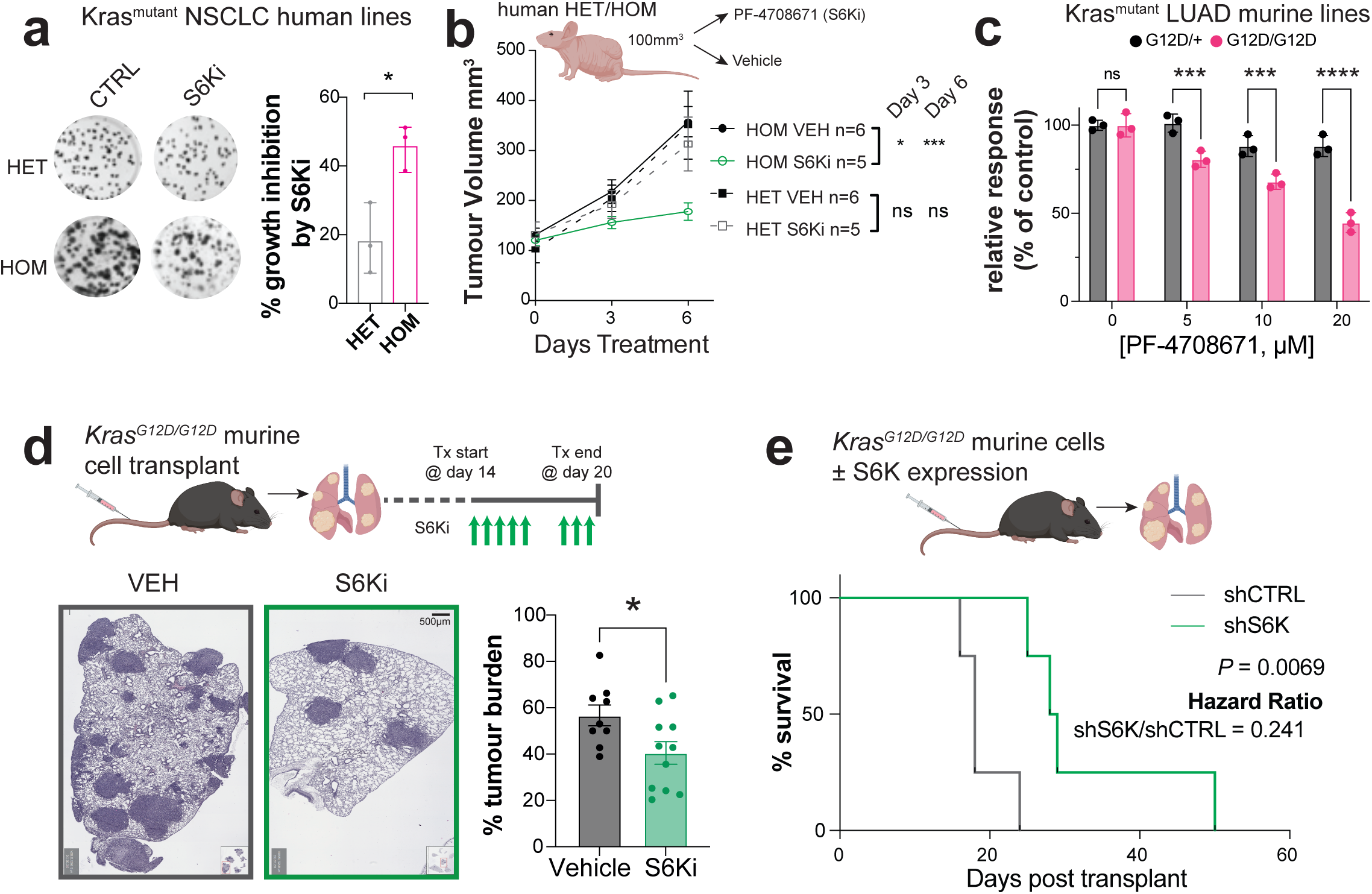
*KRAS^mutant^* copy gain drives enhanced dependency on S6K signalling. (a) Inhibition of colony formation in HET/HOM lines ± S6Ki treatment for 10 days, representative images shown, with mean % growth inhibition (1-fraction of control x 100%) ±sd indicated, analysed by t-test; (b) human HET/HOM xenograft growth ± daily S6Ki treatment, data shows cohort mean ±sem for indicated timepoints, data analysed by two-way ANOVA with Tukey post analysis testing, *p<0.05, ***p<0.001, ns = not significant; (c) response to increasing doses of PF-4708671 in murine HET/HOM cell lines after 72hrs treatment, presented as percentage of vehicle control. Data shows biological replicates ± sem, analysed by Two-way ANOVA using Šídák’s multiple comparisons testing for pairwise comparisons, ***p<0.001, ****p<0.0001; (d) Analysis of tumour burden in mice transplanted with murine HOM cell line treated with PF-4708671 as indicated. Representative images shown, and tumour burden per mouse indicated. *p<0.05 by t-test; (e) Kaplan Meier survival analysis of animals transplanted with murine HOM cells ± sh-p70S6K, significance analysed by Log-Rank testing (n=4/cohort, data representative of 2 independent experiments).

To confirm these observations we tested an independent model, the HET and HOM Kras^G12D^;p53^null^ LUAD murine cell line pair^7^ used earlier ( Ext. data figure 3d). These murine cells were overall less sensitive to PF-4708671 than human cells, nevertheless HOM cells exhibited a dose-dependent sensitivity PF-4708671 treatment in vitro, while HET cells were overtly resistant to treatment (Figure 4c). These murine HET/HOMcell lines also showed differential sensitivities to 17-AAG and AG-014699 (Ext. data figure 4c-d) identified in human in silico screen (Figure 3a) and verified in human cell panel (Ext. data figure 4a-b), confirming consistent genotype specific dependencies across human and mouse HET/HOM models. Importantly, when these HOM cells were transplanted onto immunocompetent syngeneic mice via tail vein injection, PF-4708671 significantly inhibited lung tumour growth of HOM cells, relative to vehicle treated mice (Figure 4d), confirming the anti-tumour activity of the treatment on this genotype. Murine HET cells could not be treated as they do not grow *in vivo*^7^.

To confirm that the sensitivity of HOM tumours to PF-4708671 was due to on-target activity of the inhibitor on S6K, murine HOM cells were transduced with shRNA targeting S6K (or control) to deplete p70S6K expression (Ext. data figure 6a). Again, loss of p70S6K signalling significantly reduced cell line growth *in vitro* (Ext. data Figure 6b) and *in vivo*, as indicated by significantly longer survival of animals with shS6K vs shCTRL treated HOM cells – median survival was 28 and 18 days, respectively (Figure 4e, p=0.0069).

We previously showed that HOM and HET cells have significantly distinct metabolic features^7^ with HOM cells being highly dependent on glucose to fuel glycolysis and the TCA-cycle driven production of glutathione (GSH), a fundamental metabolite for the maintenance of redox homeostasis. Given the differential sensitivity seen for PF-4708671 between HOM and HET cells, and the strong links between S6K/mTOR and glucose metabolism, we postulated that the enhanced sensitivity in HOM cells was linked to our previously reported HOM-specific glucose dependence. To test this, we used uniformly labelled ^13^C-Glucose (^13^C_6_-GLC) and examined the metabolic response to PF-4708671 treatment in HOM cells. Cells were treated for 6hrs with ^13^C_6_-GLC, before metabolites were analysed, and changes in labelling patterns were compared between CTRL and PF-4708671 groups (Figure 5a). Significant differences in glucose metabolism were observed between the treatment groups, as detailed in Ext. data Figure 7. Interestingly, we saw significant inhibition of glucose derived TCA cycle metabolites (Figure 5b, aKetoglutarate (aKG), malate) and other related metabolites (glutamate, aspartate). However, the difference in sensitivity to PF-4708671 was not simply due to a blunting of total glucose metabolic flux, as glycolysis-related metabolites increased (pyruvate, lactate). More specifically, PF-470867 treatment drove a significant drop in total GSH levels (Figure 5c) in HOM cells and in their GSH/GSSG ratio (Figure 5d), indicating that the ability of HOM cells to buffer redox stress is reduced by S6K inhibition. Together, these analyses directly link S6K signalling in MutCG cells to their unique glucose metabolism and increased dependency on glutathione production for redox buffering.

**Figure 5:**
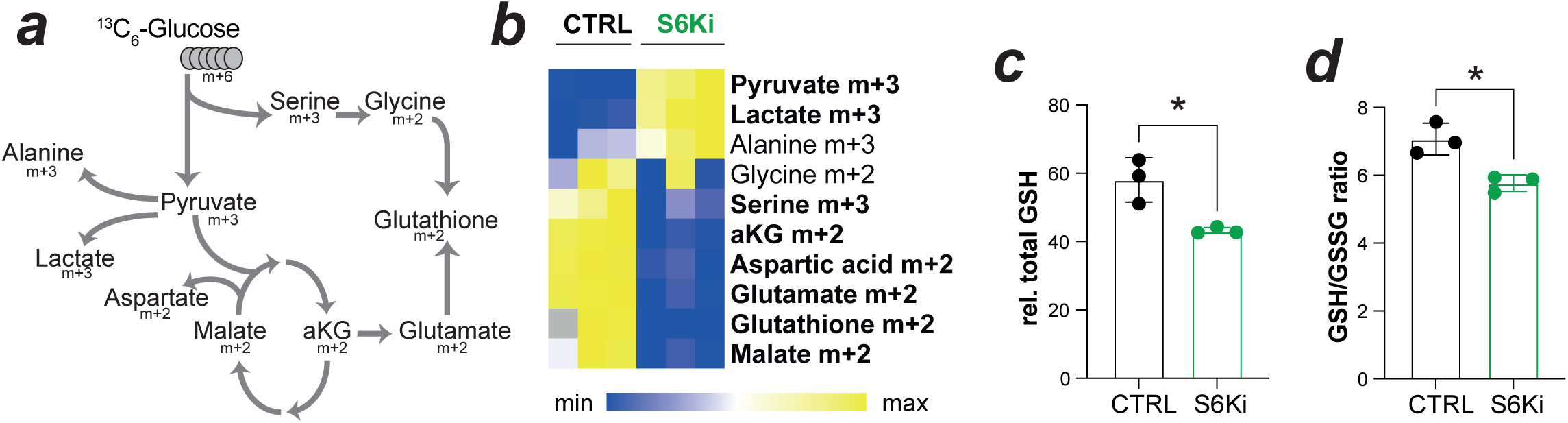
S6K inhibition blunts mitochondrial glucose metabolism and glutathione biosynthesis. (a) schematic representing labelling pattern from uniformly labelled ^13^C_6_ -glucose; (b) Relative % labelled metabolite abundance, with differentially enriched (adj. P<0.05) isotopologues indicated in bold in murine HOM cells ± S6Ki analysed by multiple t-test with Holm-Sidak correction; Relative total glutathione (c) and GSH/GSSG ratio (d) for HOM cells ± S6Ki treatment.

## Discussion

Despite remarkable improvements in clinical management and treatment, lung cancer remains a leading cause of cancer-associated mortality worldwide. Immunotherapy and an increasing number of approved targeted therapies, which now include mutant KRAS inhibitors, are undoubtedly improving depth and durability of responses. However, a high proportion of lung cancer patients, and particularly those with advanced disease, inevitably develop resistance to treatment. Multiple mechanisms of resistance to treatment have been identified^35–39^ including mutations, alterations in the tumour microenvironment and tumour hypoxia. Here, we provide evidence for a novel mechanism of resistance to chemotherapy in mutant KRAS lung cancers – mutant KRAS copy gains or amplifications.

Mutant KRAS copy gains have been identified on multiple tumour types, confirming that *KRAS^mutant^* allelic content is a frequent contributor to the genetic heterogeneity of these cancers^40–42^. Mutant gains and amplifications were also shown to be phenotypically relevant, driving invasion, metabolic rewiring and aggressive phenotypes promoting progression in NSCLC^7,8,43^, pancreatic^44,45^ and colorectal cancers^9,10^. More recently, mutant gains/amplifications were proposed as a potential mechanism of resistance to RAS inhibitors, given their enrichment in the circulating DNA of cancer patients that progressed following treatment with mutant KRAS G12C inhibitors^5,46,47^. Intriguingly, in our assays, mutant KRAS gains did not alter the sensitivity of cells to KRAS inhibitors, in agreement with previous reports^48–50^. It is plausible that in the clinic, tumour cell extrinsic mechanisms and/or previous treatments play a role in the selective advantage of cells with KRAS MutCG, upon exposure to RAS inhibitors.

Importantly, we found a new and significant correlation between KRAS gains and decreased sensitivity to cisplatin, carboplatin and ionising radiation, coupled to poorer overall survival in lung cancer. Our data demonstrate that the rewiring of glucose metabolism is likely to play a role in resistance to platinum-based chemotherapy in KRAS MutCG tumours, potentially via ROS-suppression or GSH-mediated detoxification. However, a role for alternative mechanisms of resistance cannot be fully excluded, as previously described mechanisms of cisplatin-resistance^51^ are also enriched in MutCG cells, including tetraploidy^44,52^ and altered mitochondrial apoptotic programs^53^. It is still unclear why the activity of platinum and ionising radiation are particularly affected in these cells, but not other DNA-damage inducing agents. This selectivity suggests the effect it is not linked to a global DNA-damage resistance phenotype but rather a metabolic resistance state. However, we cannot exclude that DNA repair isn’t impacted by mutant copy gains.

In depth profiling of cells with variable mutant KRAS allelic content demonstrated that mutant KRAS gains also generate actionable therapeutic liabilities, namely an increased dependency on phosphorylated p70S6K/S6 (P-S6) signalling. Indeed, cells that underwent MutCG not only expressed higher levels of P-S6, but were also significantly more sensitive to a p70S6K/S6 inhibitor than mutant heterozygous/copy neutral cells. Interestingly, this expression/sensitivity profile was specific to p70S6K/S6, as other signalling nodes in the KRAS/MAPK pathway were similarly expressed in MutCG and copy neutral mutant KRAS cells and no differential sensitivity to other pathways inhibitors was identified. Why gains in mutant KRAS copies increase dependency on p70S6K/S6 is not yet fully understood. However, increased S6 phosphorylation is a canonical readout of mTORC1-driven anabolic signalling, promoting translational capacity, cell growth and metabolic fitness – intrinsic characteristics of mutant copy gain cells. Our data shows that in KRAS MutCG cells S6 signalling helps maintain glucose carbon flow into TCA-linked metabolites and glutathione, thereby supporting redox buffering. MutCG cells gain fitness through this coordinated anabolic/stress-buffering state, and therefore become selectively dependent on the systems that maintain it. It is tempting to argue that cells with MutCG are selected for their “goldilocks phenotype”, whereby cells that upregulate only this particular axis are positively selected, as over-activation of the full RAS signalling cascade can be detrimental for their fitness^54^.

As shown here for mutant KRAS gains, enhanced S6K/mTOR signalling has been linked with more aggressive NSCLC^55–57^. Furthermore, the inhibition of the p70S6K axis specifically^29^ has demonstrated potential in controlling NSCLC growth in preclinical models^58^. However, in the clinic, targeting of this axis has given mixed results. Our data suggest that patient stratification according to KRAS status or metabolic signatures may be crucial to improve response. In agreement with our findings, Liang and colleagues reported an increase in mTOR signalling in cisplatin-resistant models of NSCLC^59^. It is thus plausible that the p70S6K axis represents a particularly attractive targeting opportunity in patients that developed resistance to platinum-based chemotherapy and show evidence of Mutant KRAS gains.

Overall, our findings here highlight the complexities associated with treating mutant *KRAS* lung cancer patients. Even in an era of precision medicine, and with mutant KRAS inhibitors becoming a very welcome reality, mutant KRAS NSCLC remains difficult to treat, with durability and depth of response lagging behind those observed with therapies targeting other known drivers. Combination therapy will likely improve responses, but a better understanding of the dependencies of mutant KRAS tumours is crucial to better inform the next generation of treatments. We show that these mutant KRAS lung cancers are genetically and metabolic heterogeneous and that *KRAS^mutant^* allelic content alone can impact lung cancer biology, patient outcome and response to therapy. Importantly, for the first time we provide a direct correlation between a measurable biomarker (*KRAS^mutant^* VAF) and therapeutic sensitivity to platinum based therapy – an important modality in NSCLC treatment; as well as targeted therapies (i.e. S6K/mTOR inhibitors). Given that KRAS MutCG are enriched in some, but not all patients treated with RAS inhibitors, we argue that allelic content heterogeneity is also likely to influence responses to future combinations of these inhibitors with chemotherapy or other agents. Our data implies that moving beyond a binary *KRAS* classification (WT/Mutant), coupled with a better understanding of the unique properties of KRAS MutCG cancers will be key for the development of curative therapies for the highly heterogenous disease typically termed as mutant KRAS NSCLC.

## Materials & Methods

### Human data

KRAS information (including variant allele frequencies, amino acid substitutions and mRNA) plus clinical information (including treatment and survival outcomes) was retrieved from cBioportal using TCGA PanCancer and Firehose Legacy samples. Binary MutCG classification was defined as VAF>0.5. Single sample scores from transcriptional analyses were generated by clara^T^ V3.0.0 platform, where calculated gene expression estimates are used within the cloud-based Almac proprietary analysis pipeline for downstream calculation of signature scores and visual reporting (Almac Diagnostic Services, https://www.almacgroup.com/diagnostics/claratreport/). Summarised Gene Expression Signature (GES) readouts (both raw and percentile ranked scores) are used for correlation to *KRAS* VAF and clinical outcome. Overall survival (OS) comparisons by Kaplan-Meier analysis using Log-rank testing for patients separated by KRAS VAF or the median gene expression score in each dataset.

### Cell lines

NSCLC cell lines H1944, H1792, H23, H358, H2030, A549, SW1573 and H460 were purchased from ATCC, and maintained in RPMI-1640 (with 10% FBS, 2mM Glutamine and 2mM Sodium Pyruvate). Murine Kras^G12D^/p53^null^ lung adenocarcinoma cell lines were isolated and characterised as previously described^7,60^, and maintained in DMEM/F12 (with 10% FBS and 2mM Glutamine). All cell lines were maintained with Penicillin/Streptomycin, tested regularly for mycoplasma and kept in a mycoplasma free environment. shRNA constructs targeting S6K or control were obtained from Origene (TR50474) and cell lines generated using standard retroviral transduction techniques.

Information on **KRAS mutation and zygosity** were retrieved from COSMIC^61^ Cell Line Project (www.cancer.sanger.ac.uk). Sanger sequencing was used to confirm mutation and zygosity on *KRAS* PCR products from genomic DNA (extracted using Qiagen DNeasy kit, #69504) using Takara Enzyme system, and primer pairs: 12/13-F: 5’TTTGAGAGCCTTTAGCCGCC3’, 12/13-R: 5’ACCCTGACATACTCCCAAGGA3’, 61-F: 5’CAGACTGTGTTTCTCCCTTC3’, 61-R: 5’GATGCAGTCTCTGGAGCAAGTT3’, following purification (QIAquick PCR purification, #28104). Chromatograms were viewed and zygosity checked using Snapgene. Pyrosequencing PCR products were analysed as previously described^7^ using Primer combinations: Codon12/13-F: 5’GGCCTGCTGAAAATGACT3’, Codon12/13-BiotR: 5’ GTTGGATCATATTCGTCCACAAA3’, Codon12/13-Seq: 5’AAACTTGTGGTAGTTGGA3’; Codon61-F: 5’CAGACTGTGTTTCTCCCTTCTCA3’, Codon61-BiotR: 5’ CCTCATGTACTGGTCCCTC3’, Codon61-Seq: TTGGATATTCTCGACACA. Samples were run on Pyromark Q24 (Qiagen) according to manufactures’ instructions, and traces analysed using Pyromark software. Genetic status of other relevant genes was compiled using cBioportal and COSMIC.

**Ras activation** assay was run as previously stated^7^, 2 x10^6^ cells were plated on 15cm culture plates (20ml RPMI 1640 media) and collected after 72 hours. Cells were pelleted, lysed and assayed using 100μg/sample (Merck Millipore, #17-497) according to manufacturer’s instructions.

For **proliferation rates**, 3x10^5^ cells were plated in triplicate on 6cm plates and live cells counted at indicated timepoints using 1:1 mix 0.4% trypan blue (Gibco) on an automated cell counter (Countess-Invitrogen, C10227). For long-term proliferation assays (colony formation), 500 cells were seeded per well and grown for 10 days before fixation and visualisation using 0.2% crystal violet. For colony formation assays with treatment, HET cell number was adjusted to 1000 cells/well, and cells were treated 24hrs after seeding for 72hrs with concentrations of drugs indicated before media change. For proliferation changes in response to drug treatment, 1x10^5^ cells per well were seeded in 6 well plates in triplicate, stained for 48-72hrs as indicated, and stained using 0.2% crystal violet or 0.05% SRB^62^. Data was normalised to vehicle control, and expressed as Fraction of Control (FoC).

**Cell viability** was assessed by Cell Titre Glo assay (Promega, #G7570, used according to manufacturer’s instructions) to screen chemo-/targeted-therapies following 72hrs treatment. IC_50_ values were calculated using non-sigmoidal dose response algorithms. SRB assay was used to screen response to KRAS inhibitors. Apoptotic cell death was analysed using Annexin V/Propidium Iodide staining, where cell pellets were resuspended in 400μL 1x binding buffer (BD #556454) containing 2μL FITC-tagged Annexin-V antibody (BD-Bioscience, excitation/emission = 485/535nm) and 3μL Propidium Iodide (Sigma-Aldrich, excitation/emission maxima of 493/636 nm). Samples were analysed by Flow cytometry, usng BD Accuri with green (FITC) and red (PI) positivity determined by FlowJo software version 10. Compensation was carried out to minimise spectral overlap. Cells were identified, gated into single cells, and death assessed using quadrant gating.

### Therapeutics

2-DeoxyGlucose (2DG, Sigma/Merck D8375) used at concentrations stated; Carboplatin (Sigma, C2538), Cisplatin (Sigma, 232120), Docetaxel (Sigma, 01885), Paclitaxel (Sigma, T7191), Vinorelbine (Cayman Chemical, 21262), Pemetrexed (Cayman Chemical, 14269), Doxorubicin (Cayman Chemical, 15007), Selumetinib (Tocris, 6815), Erlotinib (Cayman Chemical, 10483), Neratinib (Tocris, 7371), Afatinib (Cayman Chemical, 11492), MTRX849 (MedChemExpress, HY-130149), AMG-510 (Stratech, B8399-APE), MRTX-1133 (MedChemExpress, HY-134813), 17-AAG (LKT, A0025), PF-4706871 (Sigma, PZ0143), AG-014699 (SelleckChem, S1098), PD0325901 (SelleckChem, Cat.No.S1036), SCH992984 (SelleckChem, Cat.No.S7101), MK-2206 (SelleckChem, Cat.No.S1078). For dose-response assays, vehicle control was PBS or DMSO, as per compound solubility requirements.

### Radiation

Cells were irradiated with 225 kVp X-rays generated using an X-Rad 225 generator (Precision X-ray Inc., CT, USA) with a 2mm copper filter giving a half value layer (HVL) of 2.3mm Cu. All quoted doses are the absorbed dose in water 50cm from the radiation source at a dose rate of 0.591Gy/min.

**Extracellular acidification rate** (ECAR) was assessed by flux analysis (Seahorse XFe24) from technical replicates as previously described^7^, and normalised to protein content.

### Intracellular signalling

Protein expression was assessed by Western blotting using antibodies: phospho-p70S6K (#9205), total p70S6K (#2708), phospho-S6 S235 (#4858), phospho-S6 S240 (#5364), total S6 (#2217), phospho-AKT (#4060), phospho-MEK (#9154), all Cell Signalling. Equal loading was confirmed using β-actin (Sigma, A5316). Densitometry analysis preformed using Fiji software, and fold change determined as Mean relative expression HOM / Mean relative expression HET.

### In silico therapeutic analyses

IC_50_ data for 99 compounds in 29 *KRAS* mutant lung cancer cell lines was retrieved from Genomics of Drug Sensitivity in Cancer (GDSC) website (https://www.cancerrxgene.org/downloads/bulk_download, v73), and *KRAS* mutation and zygosity retrieved from COSMIC cell line project (https://cancer.sanger.ac.uk/cell_lines, filtered on KRAS/Lung). Mean IC_50_ in HOM and HET groups was determined and difference in sensitivity calculated as Mean HOM Log_ln_ IC_50_ – Mean HET Log_ln_ IC_50_. Pearson’s correlation of IC_50_ against KRAS mutant VAF in cell models tested.

### In vivo studies

Animals were maintained under SPF conditions and in compliance with UK Home Office (under PPL # P39EE17D7) or NI Department of Health (under PPL #2874) regulations. All animals were 6-8 weeks at point of transplant.

For human xenograft studies, 2x10^6^ HET (H23) and HOM (A549) cells were transplanted 1:1 in Growth Factor Reduced Matrigel (Corning, # CLS356231) into left flank of athymic nude mice and volume calculated using formula ½(a x b^2^). When tumours reached ∼100mm^3^ average per genotype, animals were randomised (n=6/group) and given PF-4708671 (50mg/kg) or vehicle (30% PEG400/ 0.5% Tween80/ 5% Propylene Glycol/PBS) daily via i.p. injection for 6 days.

For IV transplant studies, C57B/6J mice were injected with 1x10^5^ cells HOM cells (parental or shCTRL/shS6K) in PBS via lateral tail vein. Animals were recovered, observed daily, and culled at clinical endpoint of model (laboured breathing, 15% weight loss, impaired motor function, loss of appetite, dehydration). For treatments, animals were randomised (n=10 CTRL, 11 PF treated) and injected 8 times over 10 days with PF-4708671 or vehicle as above. For survival, animals were culled when the displayed >15% weight loss or reached clinical endpoint (laboured breathing, hunched posture, inactivity). For lung tumour burden, lungs were inflated, fixed overnight in 10% NBF and then paraffin embedded, before 5μm sections were stained using Haemotoxylin and Eosin.

### Statistics

Kaplan–Meier (KM) curves were carried out to compare the survival time differences. *P*-values from log-rank tests were calculated, and less than 0.05 was considered statistically significant. Data from pairwise comparisons or multiple sample/variable experiments were analysed using t-test, Fisher’s exact test, one-way ANOVA or two-way ANOVA as appropriate, with post-hoc tests as stated (Sidak, Tukey). Correlation analyses used Pearson’s algorithm. Dose response curves and IC_50_ values were calculated using non-sigmoidal algorithm. No assumptions were made of data. Laboratory data was analysed using GraphPad PRISM software.

### clara^T^ specifics

clara^T^ is a unique software-driven solution, classifying biologically relevant gene expression signatures into a comprehensive easy-to-interpret report. Version 3.0.0 clara^T^ Total mRNA Report, reports on 10 key biologies (Immuno-Oncology (IO), Epithelial to Mesenchymal Transition (EMT), Angiogenesis, Proliferation, Cell Death, Genome Instability, Energetics, Inflammation, Immortality and Evading Growth) by providing expression of 92 unique gene expression signatures, 100 single gene drug targets and 7,337 single genes relevant to the ten biologies for exploratory analysis. Visualisation of hierarchical clustering is formatted within a graphical report. Euclidean distance was calculated using the percentile ranked scores and hierarchical clustering using Ward’s linkage criteria was performed on this matrix. Samples are clustered based upon signature outputs within individual Hallmarks and across all samples.

## Contributions

Study conceptualisation and design: EMK & CPM; Human data collection, retrieval, analysis and/or interpretation: SM, GL, GJ, JB, CD, VC, NO, RK, CPM & EMK; Cell based experiments were designed, carried out and/or analysed by AB, SS, CMC, WMD, GD, DM, CPM & EMK; Metabolomics analyses were designed and carried out by AP, CB & EMK; Animal based experiments were designed, carried out and/or analysed by CMC, SM, NL, TD, AS, CPM & EMK; Manuscript prepared by CPM & EMK, with all authors contributing to editing.

## Acknowledgements

We thank the patients who contributed to all datasets used in the analyses presented. We thank the Biological Services Unit at Queen’s University Belfast and the Cambridge Biological Services at University of Cambridge for supporting the animal studies undertaken. We thank Dr Mihaela Ghita and Dr Karl Butterworth for their assistance with radiation techniques. This work was carried out with support from Lori Monroe Lung Cancer Foundation of America (to CPM) and QUB Fellowship Academy and Cancer Research UK (C61288/A26045, to EMK).

---

**Ext. Data Figure 1:**
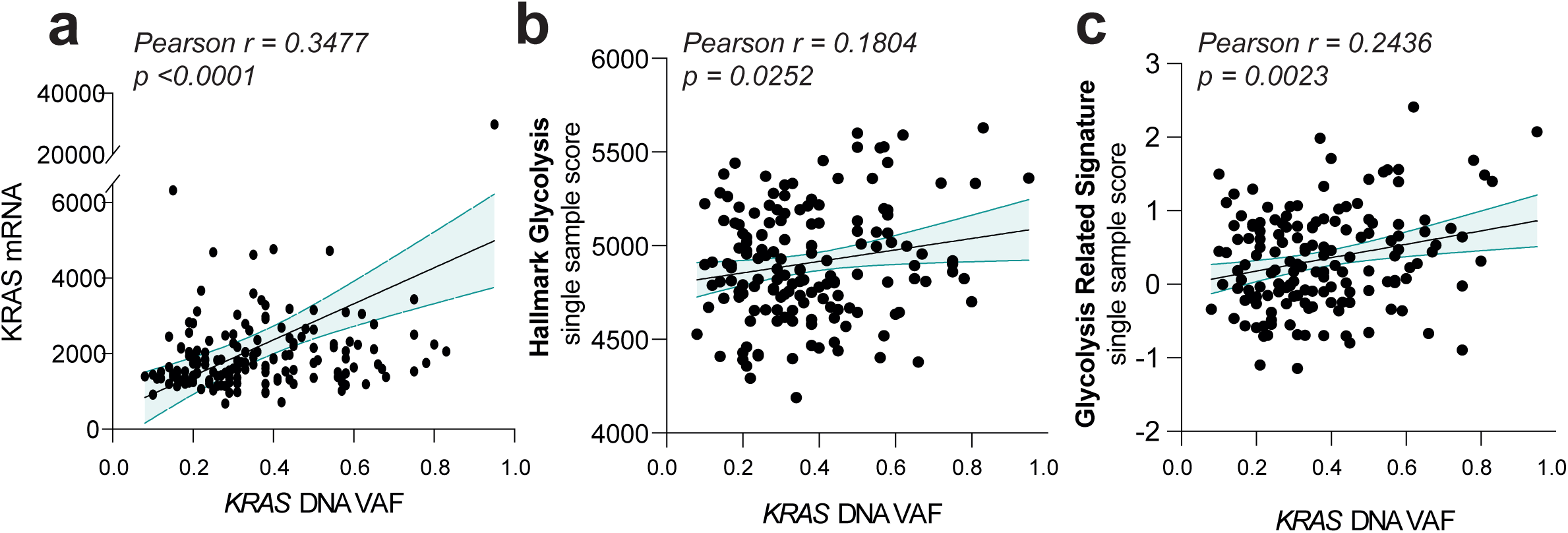
Increased glycolytic signalling driven by *KRAS^mutant^* copy gains in Lung Adenocarcinoma. (a) correlation of *KRAS^mutant^* DNA VAF vs (a) total KRAS mRNA expression, (b) MSigDB Hallmark Glycolysis score or (c) Glycolysis Related Signature score (Liu et al, 2019) per sample. Correlation by Pearson’s algorithm, with Pearson’s r and p value reported.

**Ext. Data Figure 2:**
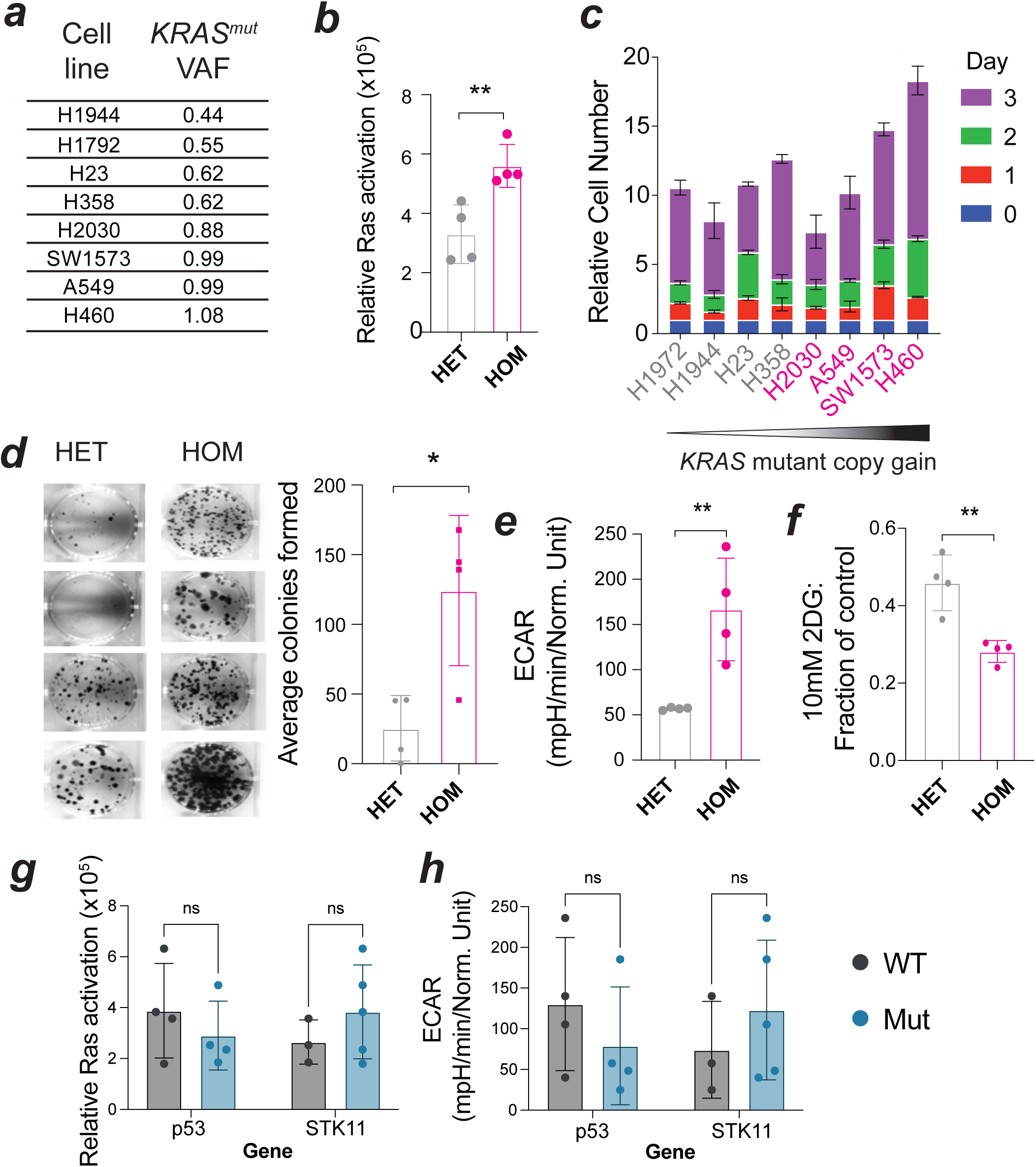
Modelling *KRAS^mutant^* copy gains in human Non-Small Cell Lung Cancer - a cell line panel for drug sensitivity testing. (a) Summary table of KRAS mutation, VAF and zygosity in 8 NSCLC cell lines; (b) Relative Ras activity in heterozygous (HET) and homozygous (HOM) cell lines, ach datapoints represent individual cell line mean of technical replicates, bar shows genotpye mean ±sem; (c) Cumulative cell growth rate per cell line over 72hrs with timepoint mean ± s.d. shown in HET (Grey) and HOM (Pink) cell lines; (d) Colony forming ability in HET and HOM cell lines, with representative image per cell line shown, graph datapoints show individual cell line means, bar indicates genotype mean ± sem; (e) Relative ECAR measured by Seahorse XFe96, normalised to crystal violet, per cell line, with bar indicating genotype mean ±sem; (f) 2DG sensitivity (shown as fraction of control, calculated from crystal violet staining of cells treated ± 10mM 2DG) per cell line, bar indicates genotype mean±sem. Relative RAS (g) and ECAR (h) in cell line panel, split as WT vs Mutant (Mut) for p53 and STK11 status. Significance determined by pairwise comparison using t-test, *p<0.05, **p<0.01, ns = not significant.

**Ext. Data Figure 3:**
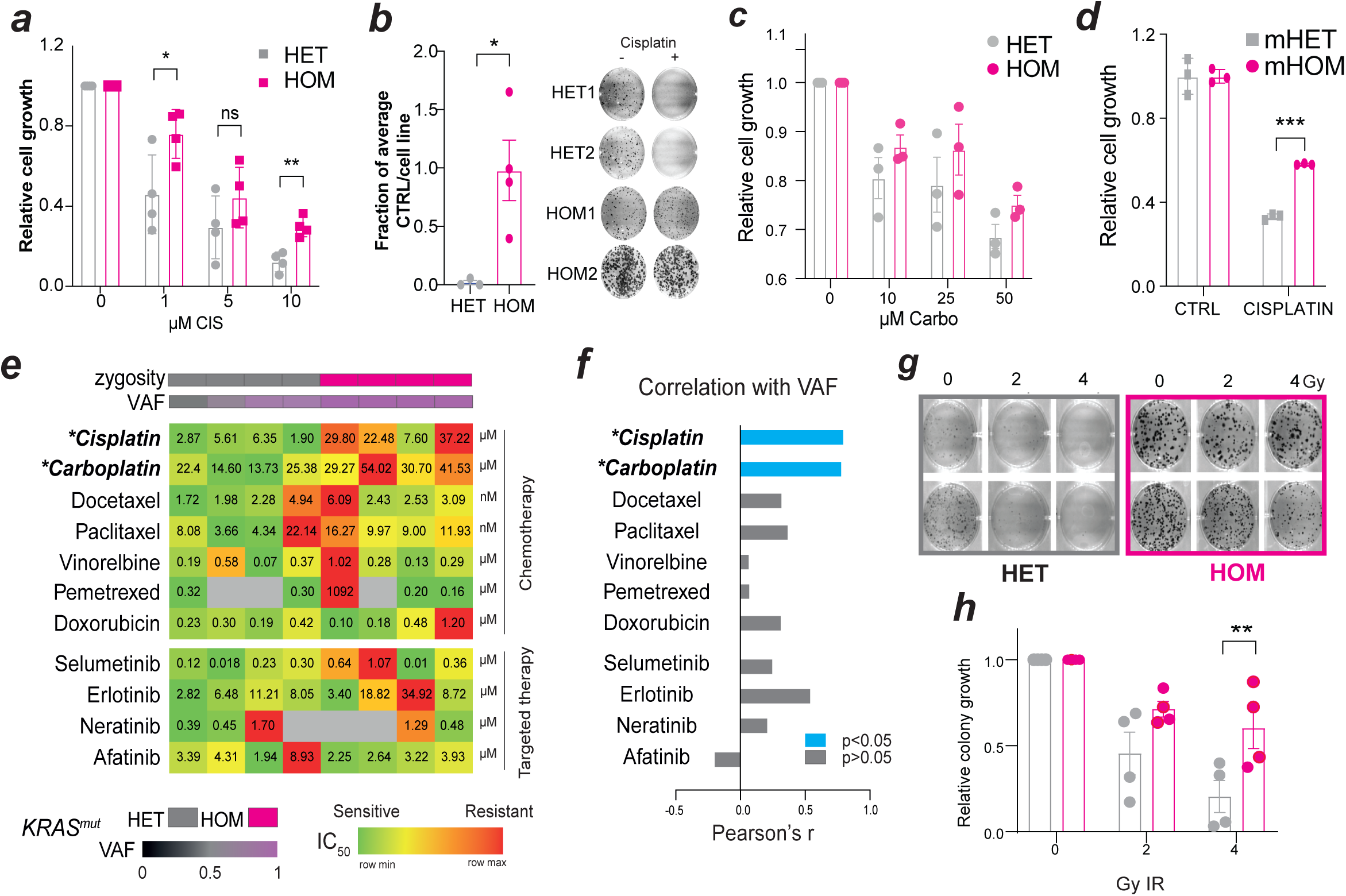
*KRAS^mutant^* homozygosity selectively promotes resistance to platinum and radiotherapy treatments. Human (a,b,d,e,f,g,h) or murine (c) *Kras^G12D^* heterozygous (HET) or homozygous (HOM) lung cancer cell lines were analysed following treatment. (a) relative cell growth in response to increasing doses of cisplatin for 72hrs determined by CV staining; (b) colony forming potential of cell lines treated with cisplatin, relative to cell line control; (c) relative cell growth in cells lines ± 10µM Cisplatin for 72hrs; (d) relative growth of cell lines ± Carboplatin at indicated doses for 72hrs; (e-f) cell lines treated with chemo/targeted therapy for 72hrs, were analysed by genotype (pairwise comparison) or against *KRAS^mutant^* VAF (correlation) as indicated. IC_50_ values for each cell line presented (e) in uM or nM as shown, red indicates higher (less sensitive) and green lower (more sensitive) cell line IC_50_ values per treatment, grey indicates IC_50_ not achieved. HOM vs HET compared by t-test, *Cisplatin *Carboplatin indicates p<0.05; (f) Pearson’s analysis of *KRAS^mutant^* VAF and IC_50_ values with r values plotted, p<0.05 represented in blue. (g-h) Colony formation assay of cell lines ± IR, colonies stained and counted after 10 days and normalised to each cell line control; Datapoints represent individual cell line mean of technical replicates and line shows genotype mean ±sem (a,b,d,h) or technical triplicates ±sd, representative of n=3 experiments (c), with representative images from n=2 lines per condition shown (b,h). Data analysed by two-way ANOVA, with Tukey’s correction (a,c,d,h) or t-test (b), *p<0.05, **p<0.01, ****p<0.0001, ns = not significant.

**Ext. data figure 4:**
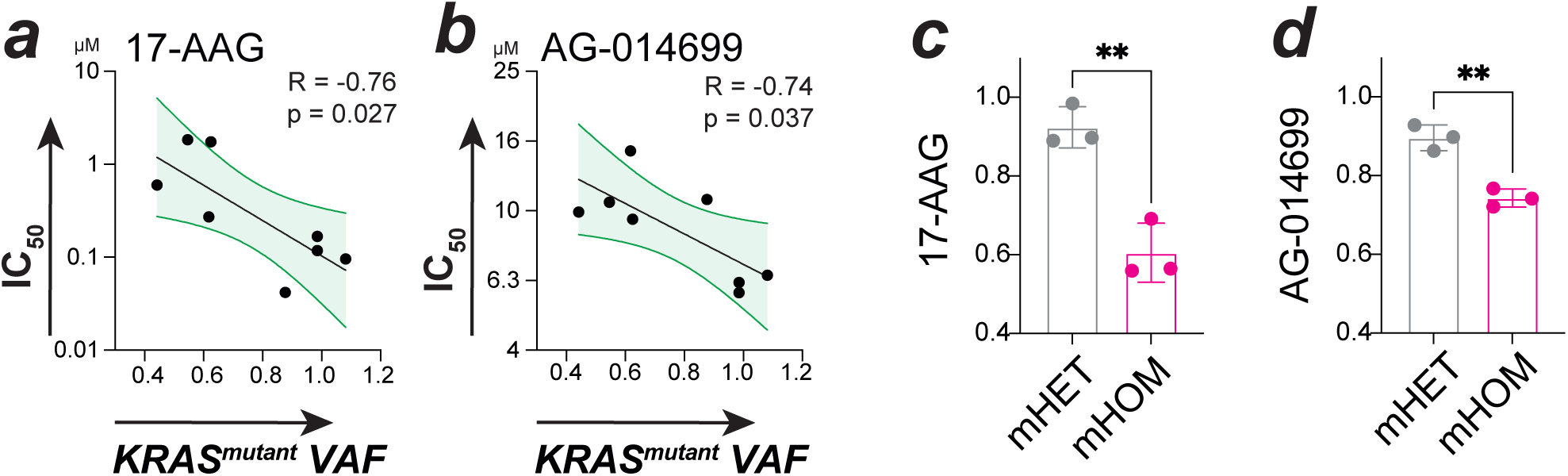
*KRAS^mutant^* copy gain promotes increased sensitivity to distinct targeted therapies. IC_50_ values correlated with *KRAS^mut^* Variant Allele Frequency (VAF) for 17-AAG (a) and AG-014699 (b) in 8 NSCLC cell line panel, with correlation analysis outputs detailed and linear regression ± 95% confidence interval shown; Relative response (fraction of each cell line control) in murine HET/HOM cell lines to same inhibitors (c-d), data shows n=3 ±s.e.m, analysed by t-test, **p<0.01.

**Ext. data figure 5:**
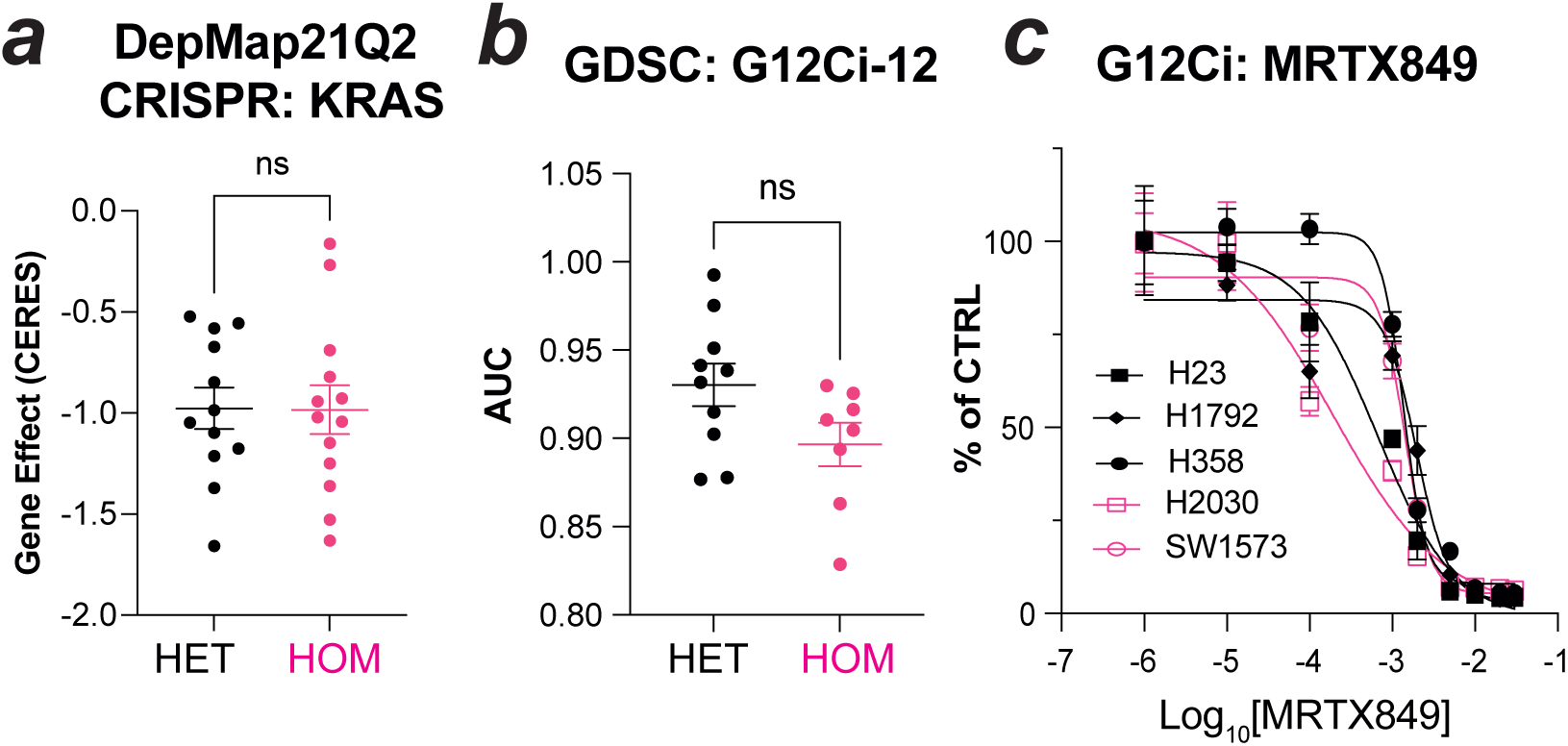
*KRAS^mutant^* copy gain does not dictate initial sensitivity to KRAS inhibition. (a) KRAS Gene Effect scores for HET and HOM NSCLC cell lines from DepMap 21Q2 CRISPR screening data, data points represent individual cell lines with genotype mean ± sem shown; (b) Genomics of Drug Sensitivity in Cancer (GDSC) screening data for KRAS G12Ci-12, data points show AUC values for HET/HOM cell lines, with genotype mean ±sem indicated; (c) MRTX849 dose response curve shown for any G12C cell lines in our panel, HET=black, HOM=pink.

**Ext. data figure 6:**
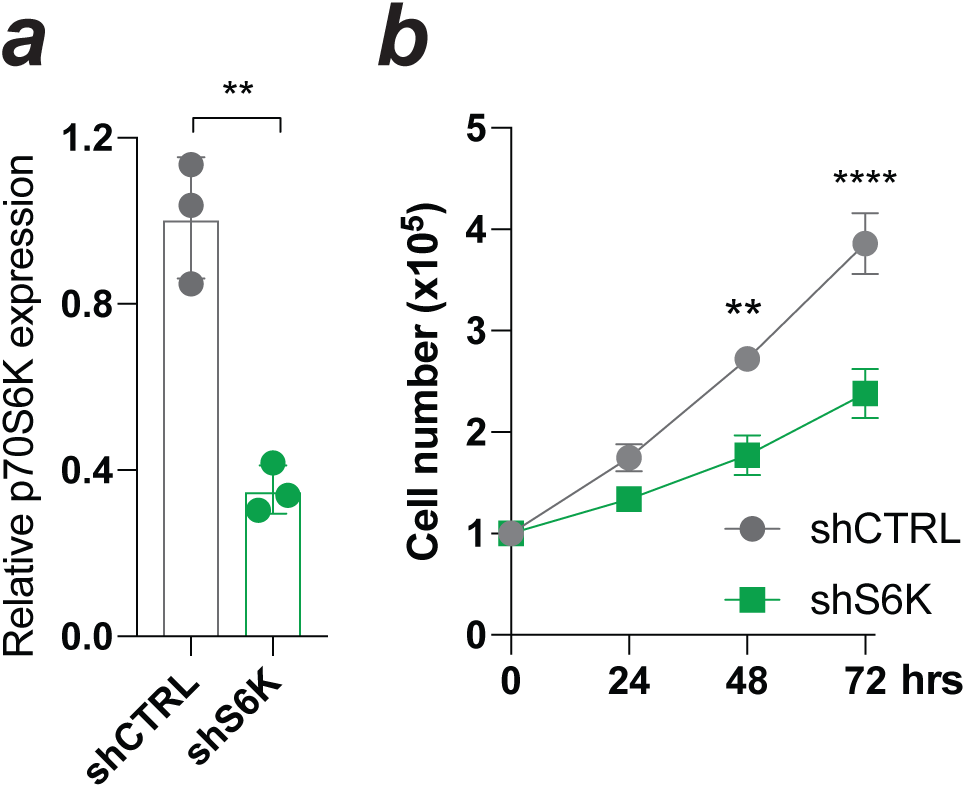
Knockdown of p70S6K impacts HOM tumour cell growth *in vitro*. (a) Relative p70S6K expression by qPCR in murine HOM lines ± sh-p70S6K, analysed by t-test; (b) Cell growth in shCTRL/shS6K murine HOM cells, data shows triplicate mean ±sd representative of n=3, analysed by multiple t-test with Holm-Šídák correction; *p<0.05, **p<0.01, ****p<0.0001.

**Ext. data figure 7:**
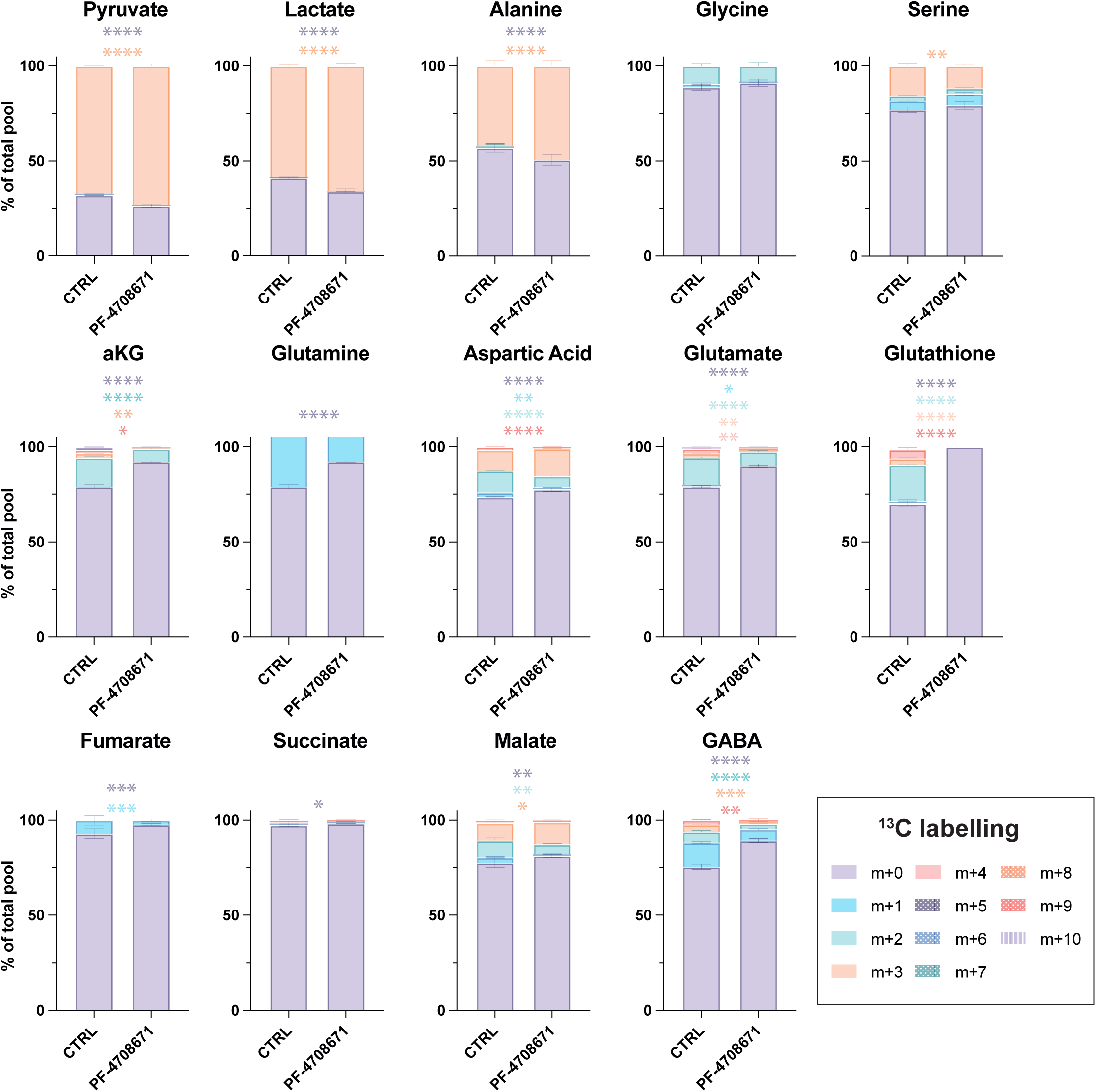
S6K inhibition blunts mitochondrial glucose metabolism. Relative % labelled isotopologue abundance following 6hr ^13^C_6_ -Glucose labelling, with significance indicated between CTRL and PF-4708671 treated groups for individual isotopologues calculated by two-way ANOVA with Šídák’s multiple comparisons test; non-significant comparisons are not shown (adj. P>0.05)

## References

1 Collisson, E. A. et al. Comprehensive molecular profiling of lung adenocarcinoma. Nature 511, 543–550 (2014). 10.1038/nature13385

2 Ma, Q., Zhang, W., Wu, K. & Shi, L. The roles of KRAS in cancer metabolism, tumor microenvironment and clinical therapy. Molecular Cancer 24, 14 (2025). 10.1186/s12943-024-02218-1

3 Stephen, Andrew G., Esposito, D., Bagni, Rachel K. & McCormick, F. Dragging Ras Back in the Ring. Cancer Cell 25, 272–281 (2014). 10.1016/j.ccr.2014.02.017

4 Cox, A. D., Fesik, S. W., Kimmelman, A. C., Luo, J. & Der, C. J. Drugging the undruggable RAS: Mission possible? Nat Rev Drug Discov 13, 828–851 (2014). 10.1038/nrd4389

5 Dilly, J. et al. Mechanisms of Resistance to Oncogenic KRAS Inhibition in Pancreatic Cancer. Cancer Discovery 14, 2135–2161 (2024). 10.1158/2159-8290.Cd-24-0177

6 Junttila, M. R. et al. Selective activation of p53-mediated tumour suppression in high-grade tumours. Nature 468, 567–571 (2010). 10.1038/nature09526

7 Kerr, E. M., Gaude, E., Turrell, F. K., Frezza, C. & Martins, C. P. Mutant Kras copy number defines metabolic reprogramming and therapeutic susceptibilities. Nature 531, 110–113 (2016). 10.1038/nature16967

8 Chiosea, S. I., Sherer, C. K., Jelic, T. & Dacic, S. KRAS mutant allele-specific imbalance in lung adenocarcinoma. Modern Pathology 24, 1571–1577 (2011). 10.1038/modpathol.2011.109

9 Burgess, M. R. et al. KRAS Allelic Imbalance Enhances Fitness and Modulates MAP Kinase Dependence in Cancer. Cell 168, 817–829.e815 (2017). 10.1016/j.cell.2017.01.020

10 Najumudeen, A. K. et al. KRAS allelic imbalance drives tumour initiation yet suppresses metastasis in colorectal cancer in vivo. Nature Communications 15, 100 (2024). 10.1038/s41467-023-44342-4

11 Yaeger, R. et al. Molecular Characterization of Acquired Resistance to KRASG12C-EGFR Inhibition in Colorectal Cancer. Cancer Discov 13, 41–55 (2023). 10.1158/2159-8290.Cd-22-0405

12 Zhao, Y. et al. Diverse alterations associated with resistance to KRAS(G12C) inhibition. Nature 599, 679–683 (2021). 10.1038/s41586-021-04065-2

13 Fennell, D. A. et al. Cisplatin in the modern era: The backbone of first-line chemotherapy for non-small cell lung cancer. Cancer Treat Rev 44, 42–50 (2016). 10.1016/j.ctrv.2016.01.003

14 Marullo, R. et al. Cisplatin induces a mitochondrial-ROS response that contributes to cytotoxicity depending on mitochondrial redox status and bioenergetic functions. PLoS One 8, e81162–e81162 (2013). 10.1371/journal.pone.0081162

15 Cerami, E. et al. The cBio Cancer Genomics Portal: An Open Platform for Exploring Multidimensional Cancer Genomics Data. Cancer Discovery 2, 401–404 (2012). 10.1158/2159-8290.Cd-12-0095

16 Gao, J. et al. Integrative analysis of complex cancer genomics and clinical profiles using the cBioPortal. Sci Signal 6, pl1 (2013). 10.1126/scisignal.2004088

17 Liu, J. et al. An Integrated TCGA Pan-Cancer Clinical Data Resource to Drive High-Quality Survival Outcome Analytics. Cell 173, 400–416.e411 (2018). 10.1016/j.cell.2018.02.052

18 Liberzon, A. et al. The Molecular Signatures Database (MSigDB) hallmark gene set collection. Cell Syst 1, 417–425 (2015). 10.1016/j.cels.2015.12.004

19 Liu, C. et al. Identification of a novel glycolysis-related gene signature that can predict the survival of patients with lung adenocarcinoma. Cell Cycle 18, 568–579 (2019). 10.1080/15384101.2019.1578146

20 Barretina, J. et al. The Cancer Cell Line Encyclopedia enables predictive modelling of anticancer drug sensitivity. Nature 483, 603–607 (2012). 10.1038/nature11003

21 Basu, A. & Krishnamurthy, S. Cellular responses to Cisplatin-induced DNA damage. J Nucleic Acids 2010 (2010). 10.4061/2010/201367

22 Siddik, Z. H. Cisplatin: mode of cytotoxic action and molecular basis of resistance. Oncogene 22, 7265–7279 (2003). 10.1038/sj.onc.1206933

23 Wangpaichitr, M. et al. The relationship of thioredoxin-1 and cisplatin resistance: its impact on ROS and oxidative metabolism in lung cancer cells. Mol Cancer Ther 11, 604–615 (2012). 10.1158/1535-7163.MCT-11-0599

24 Choi, Y.-M. et al. Mechanism of Cisplatin-Induced Cytotoxicity Is Correlated to Impaired Metabolism Due to Mitochondrial ROS Generation. PLoS One 10, e0135083 (2015). 10.1371/journal.pone.0135083

25 Chen, H. H. & Kuo, M. T. Role of glutathione in the regulation of Cisplatin resistance in cancer chemotherapy. Met Based Drugs 2010 (2010). 10.1155/2010/430939

26 De Luca, A. et al. A structure-based mechanism of cisplatin resistance mediated by glutathione transferase P1-1. Proceedings of the National Academy of Sciences 116, 13943–13951 (2019). 10.1073/pnas.1903297116

27 Yamamori, T. et al. Ionizing radiation induces mitochondrial reactive oxygen species production accompanied by upregulation of mitochondrial electron transport chain function and mitochondrial content under control of the cell cycle checkpoint. Free Radic Biol Med 53, 260–270 (2012). 10.1016/j.freeradbiomed.2012.04.033

28 Yang, W. et al. Genomics of Drug Sensitivity in Cancer (GDSC): a resource for therapeutic biomarker discovery in cancer cells. Nucleic Acids Res 41, D955–961 (2013). 10.1093/nar/gks1111

29 Pearce, Laura R. et al. Characterization of PF-4708671, a novel and highly specific inhibitor of p70 ribosomal S6 kinase (S6K1). Biochemical Journal 431, 245–255 (2010). 10.1042/bj20101024

30 Holz, M. K. & Blenis, J. Identification of S6 Kinase 1 as a Novel Mammalian Target of Rapamycin (mTOR)-phosphorylating Kinase *. Journal of Biological Chemistry 280, 26089–26093 (2005). 10.1074/jbc.M504045200

31 Magnuson, B., Ekim, B. & Fingar, D. C. Regulation and function of ribosomal protein S6 kinase (S6K) within mTOR signalling networks. Biochem J 441, 1–21 (2012). 10.1042/bj20110892

32 Fritsche-Guenther, R. et al. Alterations of mTOR signaling impact metabolic stress resistance in colorectal carcinomas with BRAF and KRAS mutations. Scientific Reports 8, 9204 (2018). 10.1038/s41598-018-27394-1

33 Byun, J.-K. et al. Oncogenic KRAS signaling activates mTORC1 through COUP-TFII-mediated lactate production. EMBO reports 20, e47451 (2019). 10.15252/embr.201847451

34 Arafeh, R., Shibue, T., Dempster, J. M., Hahn, W. C. & Vazquez, F. The present and future of the Cancer Dependency Map. Nat Rev Cancer 25, 59–73 (2025). 10.1038/s41568-024-00763-x

35 Ashrafi, A. et al. Current Landscape of Therapeutic Resistance in Lung Cancer and Promising Strategies to Overcome Resistance. Cancers (Basel) 14 (2022). 10.3390/cancers14194562

36 Chen, S. et al. Changes of tumor microenvironment in non-small cell lung cancer after TKI treatments. Front Immunol 14, 1094764 (2023). 10.3389/fimmu.2023.1094764

37 Horvath, L., Thienpont, B., Zhao, L., Wolf, D. & Pircher, A. Overcoming immunotherapy resistance in non-small cell lung cancer (NSCLC) - novel approaches and future outlook. Mol Cancer 19, 141 (2020). 10.1186/s12943-020-01260-z

38 Xu, R. et al. SIRT1/PGC-1α/PPAR-γ Correlate With Hypoxia-Induced Chemoresistance in Non-Small Cell Lung Cancer. Front Oncol 11, 682762 (2021). 10.3389/fonc.2021.682762

39 Awad, M. M. et al. Acquired Resistance to KRASG12C Inhibition in Cancer. New England Journal of Medicine 384, 2382–2393 (2021). 10.1056/NEJMoa2105281

40 Soh, J. et al. Oncogene mutations, copy number gains and mutant allele specific imbalance (MASI) frequently occur together in tumor cells. PLoS One 4, e7464 (2009). 10.1371/journal.pone.0007464

41 Yu, C. C. et al. Mutant allele specific imbalance in oncogenes with copy number alterations: Occurrence, mechanisms, and potential clinical implications. Cancer Lett 384, 86–93 (2017). 10.1016/j.canlet.2016.10.013

42 Bielski, C. M. et al. Widespread Selection for Oncogenic Mutant Allele Imbalance in Cancer. Cancer Cell 34, 852–862.e854 (2018). 10.1016/j.ccell.2018.10.003

43 Kerr, E. M. & Martins, C. P. Metabolic rewiring in mutant Kras lung cancer. Febs j 285, 28–41 (2018). 10.1111/febs.14125

44 Krasinskas, A. M., Moser, A. J., Saka, B., Adsay, N. V. & Chiosea, S. I. KRAS mutant allele-specific imbalance is associated with worse prognosis in pancreatic cancer and progression to undifferentiated carcinoma of the pancreas. Mod Pathol 26, 1346–1354 (2013). 10.1038/modpathol.2013.71

45 Yun, W.-G. et al. Multiomic quantification of the KRAS mutation dosage improves the preoperative prediction of survival and recurrence in patients with pancreatic ductal adenocarcinoma. Experimental & Molecular Medicine 57, 193–203 (2025). 10.1038/s12276-024-01382-0

46 Yaeger, R. et al. Molecular characterization of acquired resistance to KRAS G12C-EGFR inhibition in colorectal cancer. Cancer Discovery (2022). 10.1158/2159-8290.Cd-22-0405

47 Riedl, J. M. et al. Genomic landscape of clinically acquired resistance alterations in patients treated with KRAS^G12C^ inhibitors. Annals of Oncology 36, 682–692 (2025). 10.1016/j.annonc.2025.01.020

48 Chakraborty, A. et al. AZD4625 is a Potent and Selective Inhibitor of KRASG12C. Mol Cancer Ther 21, 1535–1546 (2022). 10.1158/1535-7163.Mct-22-0241

49 Canon, J. et al. The clinical KRAS(G12C) inhibitor AMG 510 drives anti-tumour immunity. Nature 575, 217–223 (2019). 10.1038/s41586-019-1694-1

50 Misale, S. et al. KRAS G12C NSCLC Models Are Sensitive to Direct Targeting of KRAS in Combination with PI3K Inhibition. Clinical Cancer Research 25, 796–807 (2019). 10.1158/1078-0432.Ccr-18-0368

51 Galluzzi, L. et al. Molecular mechanisms of cisplatin resistance. Oncogene 31, 1869–1883 (2012). 10.1038/onc.2011.384

52 Hartman, D. J., Davison, J. M., Foxwell, T. J., Nikiforova, M. N. & Chiosea, S. I. Mutant allele-specific imbalance modulates prognostic impact of KRAS mutations in colorectal adenocarcinoma and is associated with worse overall survival. Int J Cancer 131, 1810–1817 (2012). 10.1002/ijc.27461

53 Khawaja, H. et al. Bcl-xL Is a Key Mediator of Apoptosis Following KRASG12C Inhibition in KRASG12C-mutant Colorectal Cancer. Mol Cancer Ther 22, 135–149 (2023). 10.1158/1535-7163.Mct-22-0301

54 Shaw, K., Bernards, R., Stegmaier, K., Varmus, H. & Sellers, W. R. Prospects for understanding and exploiting the consequences of hyperactivation lethality. Trends in Cancer 11, 619–628 (2025). 10.1016/j.trecan.2025.04.009

55 Dhillon, T. et al. Overexpression of the mammalian target of rapamycin: a novel biomarker for poor survival in resected early stage non-small cell lung cancer. J Thorac Oncol 5, 314–319 (2010). 10.1097/JTO.0b013e3181ce6604

56 Chen, B. et al. Hyperphosphorylation of RPS6KB1, rather than overexpression, predicts worse prognosis in non-small cell lung cancer patients. PLoS One 12, e0182891 (2017). 10.1371/journal.pone.0182891

57 Zhang, Y., Ni, H. J. & Cheng, D. Y. Prognostic value of phosphorylated mTOR/RPS6KB1 in non-small cell lung cancer. Asian Pac J Cancer Prev 14, 3725–3728 (2013). 10.7314/apjcp.2013.14.6.3725

58 Qiu, Z. X., Sun, R. F., Mo, X. M. & Li, W. M. The p70S6K Specific Inhibitor PF-4708671 Impedes Non-Small Cell Lung Cancer Growth. PLoS One 11, e0147185 (2016). 10.1371/journal.pone.0147185

59 Liang, S.-Q. et al. mTOR mediates a mechanism of resistance to chemotherapy and defines a rational combination strategy to treat KRAS-mutant lung cancer. Oncogene 38, 622–636 (2019). 10.1038/s41388-018-0479-6

60 Turrell, F. K. et al. Lung tumors with distinct p53 mutations respond similarly to p53 targeted therapy but exhibit genotype-specific statin sensitivity. Genes & Development (2017). 10.1101/gad.298463.117

61 Tate, J. G. et al. COSMIC: the Catalogue Of Somatic Mutations In Cancer. Nucleic Acids Research 47, D941–D947 (2018). 10.1093/nar/gky1015

62 Vichai, V. & Kirtikara, K. Sulforhodamine B colorimetric assay for cytotoxicity screening. Nature Protocols 1, 1112–1116 (2006). 10.1038/nprot.2006.179

